# *rGREAT*: an R/Bioconductor package for functional enrichment on genomic regions

**DOI:** 10.1101/2022.06.05.494877

**Authors:** Zuguang Gu, Daniel Hübschmann

## Abstract

**Summary:** *GREAT* is a widely used tool for functional enrichment on genomic regions. However, as an online tool, it has limitations of outdated annotation data, small numbers of supported organisms and gene set collections, and not being extensible for users. Here we developed a new R/Bioconductor package named *rGREAT* which implements the *GREAT* algorithm locally. *rGREAT* by default supports more than 500 organisms and a large number of gene set collections, as well as self-provided gene sets and organisms from users. Additionally, it implements a general method for dealing with background regions.

**Availability and implementation:** The package *rGREAT* is freely available from the Bioconductor project: https://bioconductor.org/packages/rGREAT/. The development version is available at https://github.com/jokergoo/rGREAT. Gene Ontology gene sets for 556 organisms are freely available at https://jokergoo.github.io/rGREAT_genesets/.

**Supplementary information:** Supplementary data are available at Bioinformatics online.

## Introduction

Genomics and epigenomics studies often generate lists of genomic regions of interest, *e*.*g*., single-nucleotide variants (SNVs) from whole genome sequencing or exon sequencing data, peak regions of a certain chromatin modification from ChIP sequencing data, or differentially methylated regions (DMRs) from whole genome bisulfite sequencing data. The next step of analysis is naturally to associate biological functions to these genomic regions. A widely used approach is to first annotate genomic regions to the nearest genes, then to apply over-representation analysis (ORA) on the genes against a collection of gene sets where each gene set corresponds to a specific biological function. ORA is a common gene set enrichment analysis (GSEA) approach for analyzing whether a list of genes, *e*.*g*., differentially expressed genes, are enriched in a gene set (Khatri and Drăghici, 2005).

However, in the context of genomic regions, applying ORA directly on genes might not be proper. In ORA where the enrichment analysis is normally performed by Fisher’s exact test or based on hypergeometric distribution, the null assumption is that genes are independent and they have the same probability to be picked; but when dealing with genomic regions, the null assumption becomes genomic regions are uniformly distributed on the genome. Due to the fact that genes are not equally distributed on the genome and genes have different lengths, conversion from genomic regions to genes makes genes not being picked with equal probability. For example, a gene is difficult to be picked if regions are far from it, or a gene is easy to be picked if there are a cluster of regions all associated with it or the gene has a large length. These scenarios violate the null assumption of ORA and it would produce false positives and improper enrichment results on genomic regions.

The tool *GREAT* (Genomic Regions Enrichment of Annotations Tool) (McLean *et al*., 2010) was developed in 2010. The initial aim of *GREAT* was to associate biological functions to cis-regulatory elements, *e*.*g*., transcriptional factor binding sites (TFBS), but its algorithm allows it to be extended to any type of genomic regions. Instead of the gene-centric enrichment of ORA, *GREAT* converts the problem to region-centric. For a gene in a gene set, a basal domain extending its transcription start site (TSS), *e*.*g*., to upstream 5kb and downstream 1kb, is firstly established which captures the TSS-related short-range associations; next the basal domain is extended in both directions to maximal 1mb or until it reaches the neighbour gene’s basal domain, which captures the distal associations. In this way, for genes in the gene set, a list of extended TSSs are constructed and they are associated with the biological function of this gene set. Simply speaking, *GREAT* directly constructs region sets (or genomic domains) that associate with individual biological functions. The enrichment test is applied as follows. For a specific biological term in a form a of gene set, denote *p* as the fraction of its associated functional domains in the genome, *N* as the total number of input regions, *n* as the observed number of input regions that fall in the associated domains and the corresponding random variable as *X*, then *X* follows a binomial distribution *X* ∼ B(*p, N*) and the *p*-value of the enrichment is calculated as Pr(*X* ≥ *n*).

*GREAT* has been widely applied in a large number of studies, nevertheless, there are still limitations from the aspect of applications from users’ side. As an online tool, all annotation resources are only controlled by *GREAT* developers and it is not extensible by users. Current version (4.0.4) of *GREAT* only supports human and mouse, and it only supports seven gene set collections which have not yet been updated to the most recent ones. In this work, we present an R/Bioconductor package named *rGREAT*. It applies *GREAT* analysis in two ways. First it serves as a client to directly interact with the *GREAT* web service in the R environment. It automatically submits the input regions to *GREAT* and retrieves results from there. Second, it implements the *GREAT* algorithm locally and it is seamlessly integrated with the Bioconductor annotation ecosystem. On one hand, theoretically with local *GREAT*, it is possible to perform enrichment analysis on any organism and with any type of gene set collection; and on the other hand, Bioconductor annotation packages are well maintained and updated, and it ensures local *GREAT* analysis always uses the most up-to-date annotation data. Local *GREAT* by default supports many gene sets collections and more than 500 organisms, and more importantly, local *GREAT* allows providing self-defined gene sets and organisms though a simple application programming interface (API).

## Methods and results

### Online GREAT

*rGREAT* supports interacting with the *GREAT* web service programmatically. The function `submitGreatJob()` automatically submits the input regions to the *GREAT* web service, the function `getEnrichmentTables()` retrieves results from *GREAT*, and the function `plotRegionGeneAssociations()` generates plots of associations of input regions and genes that are the same as in the *GREAT* web service. `submitGreatJob()` supports all historical versions of *GREAT*.

### Local GREAT

The function `great()` implements the *GREAT* algorithm and applies *GREAT* analysis locally with a specific gene set collection on a specific organism. `great()` has integrated Gene Ontology (GO) gene sets for all supported organisms and MSigDB gene sets (Liberzon *et al*., 2011) for humans. `great()` also supports more than 500 organisms where the annotations are retrieved with the *biomaRt* package (Durinck *et al*., 2005). `great()` allows users to integrate self-provided gene sets and organisms through a simple API. The enrichment results can be viewed via a Shiny web application with the function `shinyReport()`.

### Work with background regions

*GREAT* applies a different enrichment test when background regions are provided. If denoting background regions as a list of *n* intervals (*x*_*i*_, *y*_*i*_) where the index set is *A* = {1, …, *n*}, the input regions can only be a list of intervals (*x*_*j*_, *y*_*j*_) where the index set *B* is a subset of *A*. In this setting, for a biological term, *GREAT* counts the number of background regions denoted as *N*_*bg*_, the number of input regions (or foreground regions) denoted as *N*_*fg*_, the number of background regions that fall in the associated functional domains denoted as *n*_*bg*_, the number of input regions that fall in the associated domains as *n*_*fg*_ and the corresponding random variable as *X*_*fg*_, then *X*_*fg*_ follows a hypergeometric distribution *X*_*fg*_ ∼ Hyper(*N*_*bg*_, *N*_*fg*_, *n*_*bg*_). The *p*-value is calculated as Pr(*X*_*fg*_ ≥ *n*_*fg*_).

This approach is useful in certain scenarios. For example, for a transcriptional factor (TF) whose binding sites are measured by ChIP sequencing, the union of its peaks from all tissues can be taken as the background set, and peaks from one specific tissue are taken as input region set, then we can test which biological functions are enriched for tissue-specific TFBS peaks against the background set. However, such background sets sometimes are not easy to obtain; on the other hand, researchers may look at the “background set” from a different aspect. For example, they may want to exclude assembly gap regions or unsequenced regions from the analysis. The null assumption of the *GREAT* binomial test is that input regions are uniformly distributed in the genome. Since the unsequenced regions are never measured, they are suggested to be excluded from the analysis (Domanska *et al*., 2018). In Supplementary File 1, we demonstrate indeed excluding gap regions decreases the number of enriched terms. Other scenarios of such type of backgrounds are when analyzing DMRs, the background can be set as regions showing similar CpG density as the input DMRs, or to remove sex chromsomes when genders contribute a huge batch effect in the analysis. A proper background should be selected based on the attributes of input regions, and a large and improper background normally underestimates the fractions of biological term-associated domains in the genome and it generates lower *p*-values, thus possibly more false positives (Supplementary File 1).

`great()` supports two arguments `background` and `exclude` for setting a proper background. If any of the two arguments is set, the input regions and the extended TSSs are intersected to the background, and the *GREAT* binomial algorithm is only applied to the reduced regions. When background regions are set, for a biological term, denote *N*_2_ as the total number of input regions that overlap to the background, *p*_2_ as the fraction of the associated functional domains but only in the restricted background, *n*_2_ as the observed number of input regions that fall in the associated domains restricted by background and the corresponding random variable as *X*_2_, then *X*_2_ follows the binomial distribution *X*_2_ ∼ B(*p*_2_, *N*_2_) and *p*-value is calculated as Pr(*X*_2_ ≥ *n*_2_). In fact, the native hypergeometric method in *GREAT* can be approximated to the binomial method here. Following the denotations used previously, there are *N*_2_ ≡ *N*_*fg*_, *n*_2_ ≡ *n*_*fg*_ and *p*_2_ ≈ *n*_*bg*_/*N*_*bg*_ when the numbers of regions are large. However, the binomial method is more general and it has no restriction as the hypergeometric method where input regions must be perfect subsets of backgrounds.

In Supplementary File 1, we applied functional enrichment on a TFBS dataset by taking regions in different chromatin states as backgrounds. We found TFBSs are specifically enriched in more biological functions taking enhancers as background than promoters.

### Compare online and local GREAT

In Supplementary File 2, we compared GO gene sets used in the *GREAT* web service and in local *GREAT* analysis in *rGREAT*. We found GO gene sets in *GREAT* are outdated and have inconsistencies compared to the newest ones. As a comparison, local *GREAT* always uses the newest GO annotations from the *GO*.*db* package which is updated twice a year by the Bioconductor core team. In Supplementary File 3, we also compared the enrichment results from online *GREAT* and local *GREAT* with four TFBS datasets. In general, the results from the two *GREAT* analyses are very consistent.

### Different TSS annotations

The *GREAT* method depends on the locations of TSSs. Different sources may have different annotations of TSSs. `great()` supports four sources of TSSs, which are 1. the Bioconductor TxDb packages, *e*.*g*., the *TxDb*.*Hsapiens*.*UCSC*.*hg19*.*knownGene* package for humans, 2. RefSeq genes, *e*.*g*., the RefSeq *Select* or *Curated* subset, 3. GENCODE annotations (Frankish *et al*., 2021), and 4. TSSs provided by *GREAT* itself. In Supplementary File 4, we demonstrate the four TSS sources almost cover the same set of genes in the genome, but the exact locations of TSSs differ a lot. For example, the locations of GENCODE and RefSeq TSSs have a mean difference of 1255 bp and median difference of 14bp. Nevertheless, the inconsistency of TSS locations has very little effects on the enrichment results, mainly because the difference is ignorable compared to the scales of extended TSSs. In Supplementary File 4, we demonstrate with a TFBS dataset, the enrichment results from the four TSS annotations are highly consistent.

## Conclusion

We developed a new R/Bioconductor package *rGREAT* for functional enrichment on genomic regions. *rGREAT* has integrated a large number of organisms and gene set collections. We believe it will be a useful tool for functional interpretations in genomics and epigenomics studies.

## Notes

https://github.com/jokergoo/rGREAT

### Competing Interest Statement

The authors have declared no competing interest.

