## Supplementary File 1 for "*rGREAT*: an R/Bioconductor package for functional enrichment on genomic regions": use_background.html

Use background regions


### Use background regions

###### Zuguang Gu

#### 2022-06-04

#### A general example

In this document, we will discuss the use of background regions. We first demonstrate it with a ChIP-seq TFBS dataset from UCSC table browser. Parameters are:

```
clade = Mammal
genome = Human
assembly = GRCh37/hg19
group = Regulation
track = ENCODE 3 TFBS
table: GM12878 MYB
```

We first read it as a `GRanges` object.

```
library(rGREAT)
df = read.table("data/tb_encTfChipPkENCFF215YWS_GM12878_MYB_hg19.bed")
df = df[df[, 1] %in% paste0("chr", c(1:22, "X", "Y")), ]
gr = GRanges(seqnames = df[, 1], ranges = IRanges(df[, 2] + 1, df[, 3]))
```

The next two GREAT analysis uses the whole genome as background and excludes gap regions.

```
res1 = great(gr, "GO:BP", "hg19", exclude = NULL)
res2 = great(gr, "GO:BP", "hg19", exclude = "gap")
```

And we compare the significant GO terms:

```
tb1 = getEnrichmentTable(res1)
tb2 = getEnrichmentTable(res2)

library(eulerr)
lt = list(
    genome = tb1$id[tb1$p_adjust < 0.001],
    exclude_gap = tb2$id[tb2$p_adjust < 0.001]
)
plot(euler(lt), quantities = TRUE)
```

We can see, when excluding gap regions, there are fewer significant terms left. In GREAT analysis where Binomial test is applied, denote following variables: \(N\) as total number of input regions, \(n\) as number of input regions that fall into the extended TSS regions (denote the corresponding random variable as \(X\)), and \(p\) as the fraction of extended TSS regions in the genome, then \(X \sim B(p, N)\).

When gap regions are removed from the analysis, \(N\) and \(n\) are most likely unchanged because the input regions will not overlap to the gap regions, but \(p\) gets higher because the coverage of extended TSS regions is most likely unchanged, but the background size (genome subtracting gap regions) becomes smaller. When \(p\) gets higher while \(N\) and \(n\) are unchanged, the p-value from Binomial test also gets higher (when \(p\) gets higher, the probability of obtaining \(n\) regions from \(N\) with a success rate of \(p\) increases), which decreases the number of significant terms.

The null assumption of the Binomial test is that input regions are uniformaly distributed in the genome. Since gap regions are not sequenced and input regions will never overlap to them, using whole genome as background will not be proper and it under-estimates the fraction of a function associated regions in the genome, i.e. the value of \(p\), which results more smaller p-values and possible more false positives.

This might happen for other scenations, e.g. when dealing with methylation-related regions (e.g. differentially methylated regions, DMRs), choosing a background showing similar CpG density might be more proper, which can decrease the false positives caused by the regions that will never be called as DMRs (e.g. CpG poor regions).

#### Set background by chromosomes

This scenario might be less used, but it is a good example to show the idea. In `great()`, arguments `background` and `exclude` can be set to a vector of chromosomes to keep or to remove certain chromosomes, then all analysis is restricted in the selected chromosomes. In the following example, we apply GREAT analysis only on one single chromosome at a time.

```
res_list = list()
for(chr in paste0("chr", 1:22)) {
    res_list[[chr]] = great(gr, "GO:BP", "hg19", background = chr)
}
```

Next we compare the signigicant GO terms. In the followig plot, each row is a GO term, the first heatmap shows whether the GO term is significant (red) in the corresponding chromosome and the right square heatmap shows the similarities of GO terms. GO terms are clustered into several groups by clustering the similarity matrix. Summaries of GO terms in clusters are represented as word clouds and they are attached to the heatmap.

```
sig_list = lapply(res_list, function(x) {
    tb = getEnrichmentTable(x)
    tb$id[tb$p_adjust < 0.01]
})

library(simplifyEnrichment)
simplifyGOFromMultipleLists(sig_list)
```

From the plot we can see there are some degrees of specificities of the enrichment on different chromosomes, which are mainly caused by the unequal distribution of genes on chromosomes in gene sets.

#### Set background as a set of genomic regions

It is more common to set background by a set of pre-defined genomic regions. In the following example, we use the chromatin states dataset from the same cell line as in the first example. Parameters for retrieving the data from UCSC table browser are:

```
clade = Mammal
genome = Human
assembly = GRCh37/hg19
group = Regulation
track = Broad ChromHMM
table: GM12878 ChromHMM
```

We read the chromatin states data and format it as a list where each element in the list corresponds to regions in one single chromatin state.

```
df = read.table("data/GM12878_chromHMM.bed")
df = df[df[, 1] %in% paste0("chr", c(1:22, "X", "Y")), ]
all_states = GRanges(seqnames = df[, 1], ranges = IRanges(df[, 2] + 1, df[, 3]), 
    state = gsub("^\\d+_", "", df[, 4]))
all_states = split(all_states, all_states$state)
names(all_states)
```

```
##  [1] "Active_Promoter" "Heterochrom/lo"  "Insulator"       "Poised_Promoter"
##  [5] "Repetitive/CNV"  "Repressed"       "Strong_Enhancer" "Txn_Elongation" 
##  [9] "Txn_Transition"  "Weak_Enhancer"   "Weak_Promoter"   "Weak_Txn"
```

Each time, we set background to only one chromatin state:

```
res_list = lapply(all_states, function(bg) {
    great(gr, "GO:BP", "hg19", background = bg)
})
```

We compare the numbers of significant GO terms:

```
sig_list = lapply(res_list, function(x) {
    tb = getEnrichmentTable(x)
    tb$id[tb$p_adjust < 0.05]
})
sapply(sig_list, length)
```

```
## Active_Promoter  Heterochrom/lo       Insulator Poised_Promoter  Repetitive/CNV 
##              62             317               1               0               0 
##       Repressed Strong_Enhancer  Txn_Elongation  Txn_Transition   Weak_Enhancer 
##               0             153               0               0              37 
##   Weak_Promoter        Weak_Txn 
##              18               9
```

It is interesting to see there are a lot of GO terms enriched when using heterchromatin as background. In the following parts of this document, we only look at three chromatin states: active promoter, strong enhancer and heterochromatin:

```
res_list = res_list[c("Active_Promoter", "Strong_Enhancer", "Heterochrom/lo")]
```

Next two plots demonstrate total widths of background regions in different states, and numbers of TFBS peaks falling in different background.

```
par(mar = c(4, 8, 4, 1), mfrow = c(1, 2), xpd = NA)
w = sapply(res_list, function(x) {
    sum(width(x@background))
})
barplot(w, horiz = TRUE, las = 1, xlab = "Base pairs", xlim = c(1, 2e9),
    main = "Sum of widths / background width")
n_gr = sapply(res_list, function(x) {
    x@n_total
})
barplot(n_gr, horiz = TRUE, las = 1, xlab = "Number",
    main = "Number of gr in background")
text(n_gr, 1:3, n_gr)
```

It is quite astonishing that there are only 62 TFBS peaks in heterochromatin regions which covers more than 90% of the genome. We check the top most significant GO terms:

```
tb = getEnrichmentTable(res_list[["Heterochrom/lo"]])
head(tb)
```

```
##           id                      description genome_fraction
## 1 GO:0032940                secretion by cell      0.08444135
## 2 GO:1903530  regulation of secretion by cell      0.06311058
## 3 GO:0140352                 export from cell      0.08977066
## 4 GO:0051046          regulation of secretion      0.06798635
## 5 GO:0046903                        secretion      0.09649671
## 6 GO:0010646 regulation of cell communication      0.31620709
##   observed_region_hits fold_enrichment      p_value     p_adjust mean_tss_dist
## 1                   20        3.820174 9.400148e-08 5.469682e-05        290131
## 2                   17        4.344652 1.886127e-07 5.469682e-05        284828
## 3                   20        3.593386 2.540100e-07 5.469682e-05        290131
## 4                   17        4.033068 5.371873e-07 8.675574e-05        284828
## 5                   20        3.342919 8.048350e-07 1.039847e-04        290131
## 6                   37        1.887289 4.934434e-06 4.513677e-04        367578
##   observed_gene_hits gene_set_size fold_enrichment_hyper p_value_hyper
## 1                 17           693              4.292663  3.376228e-07
## 2                 14           491              4.989507  7.140142e-07
## 3                 17           749              3.971716  9.953920e-07
## 4                 14           546              4.486901  2.496071e-06
## 5                 17           823              3.614599  3.582816e-06
## 6                 34          2806              2.120324  6.095799e-06
##   p_adjust_hyper
## 1   0.0001071705
## 2   0.0001071705
## 3   0.0001071705
## 4   0.0002015578
## 5   0.0002571665
## 6   0.0003699901
```

Take the first term “GO:0032940” for example, we can actually see, although there are only 62 TFBS peaks, 20 of them are associated with “GO:0032940” and it is almost four times higher than the expected number: 62\*0.0844 = 5.2.

However, the mean distance of “GO:0032940” associated TFBS peaks to TSS in the heterochormatin background is 290kb, and mean distance to TSS of all peaks in heterochromatin is around 350kb, which is very far from TSS. With larger distance to TSS, the more we need to be careful with the relaiblity of the associations.

```
par(mar = c(4, 8, 4, 1), xpd = NA)
dist_list = sapply(res_list, function(x) {
    gra = getRegionGeneAssociations(x)
    mean(abs(unlist(gra$dist_to_TSS)))
})
barplot(dist_list, horiz = TRUE, las = 1, xlab = "mean distance to TSS",
    main = "mean distance to TSS")
```

Therefore, we will not consider the category of heterochromatin in the analysis because although it generates many significant terms statistically, whether they make biological sense needs to be questioned.

In the end, we only compare the significant GO terms from TFBS peaks by taking promoters and enhancers as backgrounds. We use a loose cutoff for adjusted p-values (0.1) because more GO terms will give a better clustering.

```
sig_lt2 = lapply(res_list[1:2], function(x) {
    tb = getEnrichmentTable(x)
    tb$id[tb$p_adjust < 0.1]
})
simplifyGOFromMultipleLists(sig_lt2)
```
