## Supplementary File 2 for "*rGREAT*: an R/Bioconductor package for functional enrichment on genomic regions": compare_GO.html

Compare GO annotations from online and local GREAT


### Compare GO annotations from online and local GREAT

###### Zuguang Gu

#### 2022-06-04

According to the **GREAT** website, the GO gene sets they use were generated between 2011 and 2012. In this document, we are going to compare the GO gene sets integrated in GREAT web service and the GO gene sets integrated in local GREAT analysis.

**GREAT** does not provide files of gene sets they use, but in the result table from GREAT analysis, there is a column “Hyper\_Total\_Genes” which is the total number of genes in each gene set, which we can use to compare.

Total number of genes in gene sets is not affected by which input regions to use. Here we simply generate a random set of input regions.

```
library(rGREAT)
gr = randomRegions(genome = "hg19")
```

We first perform the online GREAT analysis and retrieve enrichment result for the GO:BP ontology.

```
job = submitGreatJob(gr)
tbl = getEnrichmentTables(job)
tb1 = tbl[["GO Biological Process"]]
```

Next we perform local GREAT analysis. We use the same TSS definition as online GREAT, but here GO gene sets are from the **GO.db** package (versio 3.14).

```
res2 = great(gr, "GO:BP", "GREAT:hg19", min_gene_set_size = 0)
tb2 = getEnrichmentTable(res2, min_region_hits = 0)
```

We first compare the GO terms in the two sources:

```
library(eulerr)
plot(euler(list(GREAT = tb1$ID, "GO.db" = tb2$id)), quantities = T)
```

So basically, the two GO sources have very high agreement, but **GO.db** has more additional GO terms.

The absolute numbers of GO terms are less important because some very general GO terms might be filtered out by one source. For the ease of comparison, we take the common GO terms in the two sources:

```
rownames(tb1) = tb1$ID
rownames(tb2) = tb2$id

cn = intersect(rownames(tb1), rownames(tb2))
length(cn)
```

```
## [1] 12461
```

```
tb1 = tb1[cn, ]
tb2 = tb2[cn, ]
head(tb1)
```

```
##                    ID                                                  name
## GO:0043627 GO:0043627                                  response to estrogen
## GO:0034203 GO:0034203                              glycolipid translocation
## GO:0060159 GO:0060159     regulation of dopamine receptor signaling pathway
## GO:0009636 GO:0009636                           response to toxic substance
## GO:0030216 GO:0030216                          keratinocyte differentiation
## GO:0016447 GO:0016447 somatic recombination of immunoglobulin gene segments
##            Binom_Genome_Fraction Binom_Expected Binom_Observed_Region_Hits
## GO:0043627          0.0077437310      7.7437310                         19
## GO:0034203          0.0000367138      0.0367138                          2
## GO:0060159          0.0011556260      1.1556260                          6
## GO:0009636          0.0151152000     15.1152000                         28
## GO:0030216          0.0103385700     10.3385700                         21
## GO:0016447          0.0013385960      1.3385960                          6
##            Binom_Fold_Enrichment Binom_Region_Set_Coverage Binom_Raw_PValue
## GO:0043627              2.453598                     0.019     0.0004175553
## GO:0034203             54.475430                     0.002     0.0006570551
## GO:0060159              5.191992                     0.006     0.0012299790
## GO:0009636              1.852440                     0.028     0.0017548200
## GO:0030216              2.031229                     0.021     0.0022096980
## GO:0016447              4.482308                     0.006     0.0025513850
##            Binom_Adjp_BH Hyper_Total_Genes Hyper_Expected
## GO:0043627             1                73     5.78128200
## GO:0034203             1                 1     0.07919564
## GO:0060159             1                10     0.79195640
## GO:0009636             1               210    16.63109000
## GO:0030216             1               267    21.14524000
## GO:0016447             1                22     1.74230400
##            Hyper_Observed_Gene_Hits Hyper_Fold_Enrichment
## GO:0043627                       18             3.1134960
## GO:0034203                        1            12.6269600
## GO:0060159                        5             6.3134790
## GO:0009636                       25             1.5032090
## GO:0030216                       19             0.8985475
## GO:0016447                        5             2.8697630
##            Hyper_Gene_Set_Coverage Hyper_Term_Gene_Coverage Hyper_Raw_PValue
## GO:0043627            0.0122532300               0.24657530     1.108254e-05
## GO:0034203            0.0006807352               1.00000000     7.919564e-02
## GO:0060159            0.0034036760               0.50000000     5.559241e-04
## GO:0009636            0.0170183800               0.11904760     2.655281e-02
## GO:0030216            0.0129339700               0.07116105     7.202809e-01
## GO:0016447            0.0034036760               0.22727270     2.616022e-02
##            Hyper_Adjp_BH
## GO:0043627    0.00455250
## GO:0034203    0.67073903
## GO:0060159    0.04998946
## GO:0009636    0.48279785
## GO:0030216    1.00000000
## GO:0016447    0.48207542
```

```
head(tb2)
```

```
##                    id                                           description
## GO:0043627 GO:0043627                                  response to estrogen
## GO:0034203 GO:0034203                              glycolipid translocation
## GO:0060159 GO:0060159     regulation of dopamine receptor signaling pathway
## GO:0009636 GO:0009636                           response to toxic substance
## GO:0030216 GO:0030216                          keratinocyte differentiation
## GO:0016447 GO:0016447 somatic recombination of immunoglobulin gene segments
##            genome_fraction observed_region_hits fold_enrichment      p_value
## GO:0043627    8.148643e-03                   21        2.771093 4.066589e-05
## GO:0034203    3.671436e-05                    2       58.574834 5.692329e-04
## GO:0060159    1.296032e-03                    6        4.977973 1.518087e-03
## GO:0009636    1.660764e-02                   26        1.683381 8.235280e-03
## GO:0030216    1.135782e-02                   16        1.514754 6.998219e-02
## GO:0016447    3.910585e-03                    6        1.649782 1.606168e-01
##             p_adjust mean_tss_dist observed_gene_hits gene_set_size
## GO:0043627 0.6518335        186020                 19            74
## GO:0034203 1.0000000         45530                  1             1
## GO:0060159 1.0000000        250093                  5            11
## GO:0009636 1.0000000        179767                 24           246
## GO:0030216 1.0000000        454506                 13           137
## GO:0016447 1.0000000        133789                  5            54
##            fold_enrichment_hyper p_value_hyper p_adjust_hyper
## GO:0043627              3.229414  0.0000035320    0.001887147
## GO:0034203             12.577717  0.0795056868    0.652198902
## GO:0060159              5.717144  0.0009699808    0.071980661
## GO:0009636              1.227094  0.1734896379    0.957598280
## GO:0030216              1.193506  0.2939413537    1.000000000
## GO:0016447              1.164603  0.4306255491    1.000000000
```

The column `"Hyper_Total_Genes"` in `tb1` and the column `"gene_set_size"` in `tb2` all correspond to the numbers of genes in GO gene sets. We can directly compare the two columns of values.

```
plot(tb1$Hyper_Total_Genes, tb2$gene_set_size,
    xlab = "online-GREAT", ylab = "GO.db", main = "Gene set sizes")
```

In general, the two vectors agrees very linearly.

Next we add a third source of GO gene sets, which is from MSigDB. Similarly, we perform local GREAT analysis and extract the enrichment table.

```
res3 = great(gr, "msigdb:C5:GO:BP", "GREAT:hg19", min_gene_set_size = 0)
tb3 = getEnrichmentTable(res3, min_region_hits = 0)
head(tb3)
```

```
##                                                            id genome_fraction
## 1                                   GOBP_RESPONSE_TO_ESTROGEN    0.0080626915
## 2            GOBP_NEGATIVE_REGULATION_OF_PROTEIN_LOCALIZATION    0.0206171662
## 3      GOBP_REGULATION_OF_DOPAMINE_RECEPTOR_SIGNALING_PATHWAY    0.0012960321
## 4                     GOBP_XENOBIOTIC_TRANSMEMBRANE_TRANSPORT    0.0005443128
## 5                GOBP_NEGATIVE_REGULATION_OF_TELOMERE_CAPPING    0.0005762874
## 6 GOBP_PROTECTION_FROM_NON_HOMOLOGOUS_END_JOINING_AT_TELOMERE    0.0006031903
##   observed_region_hits fold_enrichment      p_value  p_adjust mean_tss_dist
## 1                   20        2.667270 0.0001017811 0.7794394        232702
## 2                   35        1.825392 0.0006473291 1.0000000        203768
## 3                    6        4.977973 0.0015180868 1.0000000        250093
## 4                    4        7.901845 0.0018224919 1.0000000         72246
## 5                    4        7.463420 0.0022372901 1.0000000        459899
## 6                    4        7.130545 0.0026332297 1.0000000        483791
##   observed_gene_hits gene_set_size fold_enrichment_hyper p_value_hyper
## 1                 18            67              3.379088  3.147781e-06
## 2                 32           189              2.129560  3.616861e-05
## 3                  5            11              5.717144  9.699808e-04
## 4                  4             7              7.187267  1.145139e-03
## 5                  2             8              3.144429  1.283857e-01
## 6                  2            10              2.515543  1.860289e-01
##   p_adjust_hyper
## 1    0.001084849
## 2    0.005501328
## 3    0.048869166
## 4    0.053058302
## 5    0.660882238
## 6    0.772062887
```

In MSigDB, the IDs of GO gene sets are GO term names, thus we need to convert them to GO IDs:

```
library(GO.db)
lt = as.list(GOTERM)
map = sapply(lt, function(x) Term(x))
map = map[sapply(lt, function(x) Ontology(x) == "BP")]
map = toupper(map)
map = gsub(" ", "_", map)
map2 = structure(names(map), names = map)
new_rn = map2[ gsub("^GOBP_", "", tb3$id) ]
l = !is.na(new_rn)
tb3 = tb3[l, ]
rownames(tb3) = new_rn[l]
```

Next we take the common GO terms in the three sources:

```
cn = intersect(cn, rownames(tb3))
length(cn)
```

```
## [1] 5625
```

```
tb1 = tb1[cn, ]
tb2 = tb2[cn, ]
tb3 = tb3[cn, ]
```

And we compare the gene set sizes in the three sources:

```
par(mfrow = c(1, 3))
max = max(tb1$Hyper_Total_Genes, tb2$gene_set_size, tb3$gene_set_size)
plot(tb1$Hyper_Total_Genes, tb2$gene_set_size, xlim = c(0, max), ylim = c(0, max),
    xlab = "online-GREAT", ylab = "GO.db", main = "Gene set sizes")
plot(tb1$Hyper_Total_Genes, tb3$gene_set_size, xlim = c(0, max), ylim = c(0, max),
    xlab = "online-GREAT", ylab = "MSigDB", main = "Gene set sizes")
plot(tb2$gene_set_size, tb3$gene_set_size, xlim = c(0, max), ylim = c(0, max),
    xlab = "GO.db", ylab = "MSigDB", main = "Gene set sizes")
```

So here we see GO gene sets from **GO.db** and MSigDb are almost identical (the third plot), while there are certain degrees of inconsistency between online GREAT and the other two (the first two plots).

Now we can make the conclusion that the GO gene sets in online GREAT are outdated and are not consistent to the most up-to-date ones.
