## Supplementary File 2 for "*rGREAT*: an R/Bioconductor package for functional enrichment on genomic regions": compare_GO.Rmd

````
---
title: "Compare GO annotations from online and local GREAT"
author: "Zuguang Gu"
date: '`r Sys.Date()`'
output: html_document
---

```{r, echo = FALSE, message = FALSE}
library(knitr)
knitr::opts_chunk$set(
    error = FALSE,
    tidy  = FALSE,
    message = FALSE,
    fig.align = "center")
```

[According to the **GREAT** website](https://great-help.atlassian.net/wiki/spaces/GREAT/pages/655448/Gene+Ontology),
the GO gene sets they use were generated between 2011 and 2012. In this document, we are going to compare the GO gene sets integrated
in GREAT web service and the GO gene sets integrated in local GREAT analysis.

**GREAT** does not provide files of gene sets they use, but in the result table from GREAT analysis,
there is a column "Hyper_Total_Genes" which is the total number of genes in each gene set, which we can use to compare.

Total number of genes in gene sets is not affected by which input regions to use. Here we simply generate a random set
of input regions.

```{r}
library(rGREAT)
gr = randomRegions(genome = "hg19")
```


We first perform the online GREAT analysis and retrieve enrichment result for the GO:BP ontology.


```{r}
job = submitGreatJob(gr)
tbl = getEnrichmentTables(job)
tb1 = tbl[["GO Biological Process"]]
```

Next we perform local GREAT analysis. We use the same TSS definition as online GREAT, but here GO gene sets are from the **GO.db** package (versio 3.14).

```{r}
res2 = great(gr, "GO:BP", "GREAT:hg19", min_gene_set_size = 0)
tb2 = getEnrichmentTable(res2, min_region_hits = 0)
```

We first compare the GO terms in the two sources:

```{r}
library(eulerr)
plot(euler(list(GREAT = tb1$ID, "GO.db" = tb2$id)), quantities = T)
```

So basically, the two GO sources have very high agreement, but **GO.db** has more additional GO terms.

The absolute numbers of GO terms are less important because some very general GO terms might be filtered out by one source.
For the ease of comparison, we take the common GO terms in the two sources:

```{r}
rownames(tb1) = tb1$ID
rownames(tb2) = tb2$id

cn = intersect(rownames(tb1), rownames(tb2))
length(cn)
tb1 = tb1[cn, ]
tb2 = tb2[cn, ]
head(tb1)
head(tb2)
```

The column `"Hyper_Total_Genes"` in `tb1` and the column `"gene_set_size"` in `tb2` all correspond
to the numbers of genes in GO gene sets. We can directly compare the two columns of values.

```{r, fig.width = 5, fig.height =5}
plot(tb1$Hyper_Total_Genes, tb2$gene_set_size,
    xlab = "online-GREAT", ylab = "GO.db", main = "Gene set sizes")
```

In general, the two vectors agrees very linearly.

Next we add a third source of GO gene sets, which is from MSigDB. Similarly,
we perform local GREAT analysis and extract the enrichment table.

```{r}
res3 = great(gr, "msigdb:C5:GO:BP", "GREAT:hg19", min_gene_set_size = 0)
tb3 = getEnrichmentTable(res3, min_region_hits = 0)
head(tb3)
```

In MSigDB, the IDs of GO gene sets are GO term names, thus we need to convert them to GO IDs:

```{r}
library(GO.db)
lt = as.list(GOTERM)
map = sapply(lt, function(x) Term(x))
map = map[sapply(lt, function(x) Ontology(x) == "BP")]
map = toupper(map)
map = gsub(" ", "_", map)
map2 = structure(names(map), names = map)
new_rn = map2[ gsub("^GOBP_", "", tb3$id) ]
l = !is.na(new_rn)
tb3 = tb3[l, ]
rownames(tb3) = new_rn[l]
```

Next we take the common GO terms in the three sources:

```{r}
cn = intersect(cn, rownames(tb3))
length(cn)
tb1 = tb1[cn, ]
tb2 = tb2[cn, ]
tb3 = tb3[cn, ]
```

And we compare the gene set sizes in the three sources:


```{r, fig.width = 10, fig.height = 10/3}
par(mfrow = c(1, 3))
max = max(tb1$Hyper_Total_Genes, tb2$gene_set_size, tb3$gene_set_size)
plot(tb1$Hyper_Total_Genes, tb2$gene_set_size, xlim = c(0, max), ylim = c(0, max),
    xlab = "online-GREAT", ylab = "GO.db", main = "Gene set sizes")
plot(tb1$Hyper_Total_Genes, tb3$gene_set_size, xlim = c(0, max), ylim = c(0, max),
    xlab = "online-GREAT", ylab = "MSigDB", main = "Gene set sizes")
plot(tb2$gene_set_size, tb3$gene_set_size, xlim = c(0, max), ylim = c(0, max),
    xlab = "GO.db", ylab = "MSigDB", main = "Gene set sizes")
```


So here we see GO gene sets from **GO.db** and MSigDb are almost identical (the third plot), while
there are certain degrees of inconsistency between online GREAT and the other two (the first two plots).

Now we can make the conclusion that the GO gene sets in online GREAT are outdated and are not consistent
to the most up-to-date ones.
````
