## Supplementary File 3 for "*rGREAT*: an R/Bioconductor package for functional enrichment on genomic regions": compare_online_and_local.html

Compare online and local GREAT


### Compare online and local GREAT

###### Zuguang Gu

#### 2022-06-04

```
library(rGREAT)
library(eulerr)
```

In this document, we will compare the enrichment results from online GREAT and local GREAT. The four datasets are all from UCSC table browser. Parameters are:

```
clade = Mammal
genome = Human
assembly = GRCh37/hg19
group = Regulation
track = ENCODE 3 TFBS
table: A549 ATF3, A549 ELF1, H1-hESC RXRA, GM12878 MYB
```

We first read the files into `GRanges` objects:

```
read_bed = function(f) {
    df = read.table(f)
    df = df[df[, 1] %in% paste0("chr", c(1:22, "X", "Y")), ]
    GRanges(seqnames = df[, 1], ranges = IRanges(df[, 2] + 1, df[, 3]))
}
grl = list()
grl$A549_JUN = read_bed("data/tb_encTfChipPkENCFF708LCH_A549_JUN_hg19.bed")
grl$A549_ELF1 = read_bed("data/tb_encTfChipPkENCFF533NIV_A549_ELF1_hg19.bed")
grl$H1_hESC_RXRA = read_bed("data/tb_encTfChipPkENCFF369JAI_H1_hESC_RXRA_hg19.bed")
grl$GM12878_MYB = read_bed("data/tb_encTfChipPkENCFF215YWS_GM12878_MYB_hg19.bed")
sapply(grl, length)
```

```
##     A549_JUN    A549_ELF1 H1_hESC_RXRA  GM12878_MYB 
##         1726        11577         2092         3748
```

#### A549\_JUN (1726 input regions)

Apply both online and local GREAT analysis. Note online GREAT exclude gap regions, thus in local GREAT, gap regions are removed as well.

```
gr = grl$A549_JUN
job = submitGreatJob(gr)
tbl = getEnrichmentTables(job)
tb1 = tbl[["GO Biological Process"]]

res = great(gr, "GO:BP", "hg19")
tb2 = getEnrichmentTable(res)
```

Overlap of significant GO terms, but only those GO terms both existing in the two enrichment tables.

```
cn = intersect(tb1$ID, tb2$id)
length(cn)
```

```
## [1] 4533
```

```
rownames(tb1) = tb1$ID
rownames(tb2) = tb2$id
tb1 = tb1[cn, ]
tb2 = tb2[cn, ]
```

```
lt2 = list(online = tb1$ID[tb1$Binom_Adjp_BH < 0.001],
          local = tb2$id[tb2$p_adjust < 0.001])
plot(euler(lt2), quantities = TRUE, main = "A549_JUN")
```

Next we compare the observed region hits and fold enrichment in the two results.

```
par(mfrow = c(1, 2))
plot(tb1$Binom_Observed_Region_Hits, tb2$observed_region_hits, pch = 16, col = "#00000010",
    xlab = "online GREAT", ylab = "local GREAT", main = "Observed region hits")
plot(tb1$Binom_Fold_Enrichment, tb2$fold_enrichment, pch = 16, col = "#00000010",
    xlab = "online GREAT", ylab = "local GREAT", main = "Fold enrichment")
```

Next we compare the two significant GO term lists by clustering them into groups.

```
lt3 = list(online = data.frame(id = tb1$ID, p_adjust = tb1$Binom_Adjp_BH), 
           local = data.frame(id = tb2$id, p_adjust = tb2$p_adjust))
library(simplifyEnrichment)
se_opt$verbose = FALSE
simplifyGOFromMultipleLists(lt3, padj_cutoff = 0.001)
```

#### A549\_ELF1 (11577 input regions)

Apply both online and local GREAT analysis:

```
gr = grl$A549_ELF1
job = submitGreatJob(gr)
tbl = getEnrichmentTables(job)
tb1 = tbl[["GO Biological Process"]]

res = great(gr, "GO:BP", "hg19")
tb2 = getEnrichmentTable(res)
```

Overlap of significant GO terms, but only those GO terms both existing in the two enrichment tables.

```
cn = intersect(tb1$ID, tb2$id)
length(cn)
```

```
## [1] 7532
```

```
rownames(tb1) = tb1$ID
rownames(tb2) = tb2$id
tb1 = tb1[cn, ]
tb2 = tb2[cn, ]
```

```
lt2 = list(online = tb1$ID[tb1$Binom_Adjp_BH < 0.001],
          local = tb2$id[tb2$p_adjust < 0.001])
plot(euler(lt2), quantities = TRUE, main = "A549_ELF1")
```

Next we compare the observed region hits and fold enrichment in the two results.

```
par(mfrow = c(1, 2))
plot(tb1$Binom_Observed_Region_Hits, tb2$observed_region_hits, pch = 16, col = "#00000010",
    xlab = "online GREAT", ylab = "local GREAT", main = "Observed region hits")
plot(tb1$Binom_Fold_Enrichment, tb2$fold_enrichment, pch = 16, col = "#00000010",
    xlab = "online GREAT", ylab = "local GREAT", main = "Fold enrichment")
```

Next we compare the two significant GO term lists by clustering them into groups.

```
lt3 = list(online = data.frame(id = tb1$ID, p_adjust = tb1$Binom_Adjp_BH), 
           local = data.frame(id = tb2$id, p_adjust = tb2$p_adjust))
library(simplifyEnrichment)
se_opt$verbose = FALSE
simplifyGOFromMultipleLists(lt3, padj_cutoff = 0.001)
```

#### H1\_hESC\_RXRA (2092 input regions)

Apply both online and local GREAT analysis:

```
gr = grl$H1_hESC_RXRA
job = submitGreatJob(gr)
tbl = getEnrichmentTables(job)
tb1 = tbl[["GO Biological Process"]]

res = great(gr, "GO:BP", "hg19")
tb2 = getEnrichmentTable(res)
```

Overlap of significant GO terms, but only those GO terms both existing in the two enrichment tables.

```
cn = intersect(tb1$ID, tb2$id)
length(cn)
```

```
## [1] 4686
```

```
rownames(tb1) = tb1$ID
rownames(tb2) = tb2$id
tb1 = tb1[cn, ]
tb2 = tb2[cn, ]
```

```
lt2 = list(online = tb1$ID[tb1$Binom_Adjp_BH < 0.001],
          local = tb2$id[tb2$p_adjust < 0.001])
plot(euler(lt2), quantities = TRUE, main = "H1_hESC_RXRA")
```

Next we compare the observed region hits and fold enrichment in the two results.

```
par(mfrow = c(1, 2))
plot(tb1$Binom_Observed_Region_Hits, tb2$observed_region_hits, pch = 16, col = "#00000010",
    xlab = "online GREAT", ylab = "local GREAT", main = "Observed region hits")
plot(tb1$Binom_Fold_Enrichment, tb2$fold_enrichment, pch = 16, col = "#00000010",
    xlab = "online GREAT", ylab = "local GREAT", main = "Fold enrichment")
```

Next we compare the two significant GO term lists by clustering them into groups.

```
lt3 = list(online = data.frame(id = tb1$ID, p_adjust = tb1$Binom_Adjp_BH), 
           local = data.frame(id = tb2$id, p_adjust = tb2$p_adjust))
library(simplifyEnrichment)
se_opt$verbose = FALSE
simplifyGOFromMultipleLists(lt3, padj_cutoff = 0.001)
```

#### GM12878\_MYB (3748 input regions)

Apply both online and local GREAT analysis:

```
gr = grl$GM12878_MYB
job = submitGreatJob(gr)
tbl = getEnrichmentTables(job)
tb1 = tbl[["GO Biological Process"]]

res = great(gr, "GO:BP", "hg19")
tb2 = getEnrichmentTable(res)
```

Overlap of significant GO terms, but only those GO terms both existing in the two enrichment tables.

```
cn = intersect(tb1$ID, tb2$id)
length(cn)
```

```
## [1] 5723
```

```
rownames(tb1) = tb1$ID
rownames(tb2) = tb2$id
tb1 = tb1[cn, ]
tb2 = tb2[cn, ]
```

```
lt2 = list(online = tb1$ID[tb1$Binom_Adjp_BH < 0.001],
          local = tb2$id[tb2$p_adjust < 0.001])
plot(euler(lt2), quantities = TRUE, main = "GM12878_MYB")
```

Next we compare the observed region hits and fold enrichment in the two results.

```
par(mfrow = c(1, 2))
plot(tb1$Binom_Observed_Region_Hits, tb2$observed_region_hits, pch = 16, col = "#00000010",
    xlab = "online GREAT", ylab = "local GREAT", main = "Observed region hits")
plot(tb1$Binom_Fold_Enrichment, tb2$fold_enrichment, pch = 16, col = "#00000010",
    xlab = "online GREAT", ylab = "local GREAT", main = "Fold enrichment")
```

Next we compare the two significant GO term lists by clustering them into groups.

```
lt3 = list(online = data.frame(id = tb1$ID, p_adjust = tb1$Binom_Adjp_BH), 
           local = data.frame(id = tb2$id, p_adjust = tb2$p_adjust))
library(simplifyEnrichment)
se_opt$verbose = FALSE
simplifyGOFromMultipleLists(lt3, padj_cutoff = 0.001)
```
