## Supplementary File 4 for "*rGREAT*: an R/Bioconductor package for functional enrichment on genomic regions": compare_tss.html

Compare different TSS sources


### Compare different TSS sources

###### Zuguang Gu

#### 2022-06-04

`great()` supports various sources to obtain gene TSS. The sources can be one of:

- a `TxDb.*` package such as **TxDb.Hsapiens.UCSC.hg38.knownGene**,
- RefSeq database (mainly the “RefSeq Select” subset (see https://genome.ucsc.edu/cgi-bin/hgTrackUi?db=hg38&g=refSeqComposite for different subsets of RefSeq),
- Gencode annotation,
- TSS provided by GREAT itself.

In this document, we will compare different TSS sources and their influence on GREAT enrichment analysis.

We use human genome hg38 here because there will be gene ID conversions (e.g. from Ensembl ID to Entrez ID for Gencode annotation), using the newest genome annotation version will reduce the inconsistency between different sources.

The helper function `getTSS()` extracts TSS from a specific source. Note genes in the first four sources are protein-coding genes.

```
library(rGREAT)

tss_txdb = getTSS("TxDb.Hsapiens.UCSC.hg38.knownGene")
tss_gencode = getTSS("gencode_v40")
tss_refseq = getTSS("refseq:hg38")
tss_great = getTSS("great:hg38")
```

Gene IDs in `tss_gencode` are Ensembl gene IDs, and gene IDs in `tss_great` are gene symbols. We convert them to Entrez gene IDs.

```
library(org.Hs.eg.db)

map = unlist(as.list(org.Hs.egENSEMBL2EG))
new_gene_id = map[tss_gencode$gene_id]
tss_gencode$gene_id[!is.na(new_gene_id)] = new_gene_id[!is.na(new_gene_id)]

map = unlist(as.list(org.Hs.egSYMBOL2EG))
new_gene_id = map[tss_great$gene_id]
tss_great$gene_id[!is.na(new_gene_id)] = new_gene_id[!is.na(new_gene_id)]
```

We put all TSS objects into a single list:

```
tss_lt = list(
    txdb_known_gene = tss_txdb,
    gencode = tss_gencode,
    refseq = tss_refseq,
    great = tss_great
)
tss_lt = lapply(tss_lt, sort)
```

We first look at the overlap of genes. It basically shows all five sources almost contain the same set of genes.

```
library(ComplexHeatmap)
lt = lapply(tss_lt, function(x) {
    unique(x$gene_id)
})
cm = make_comb_mat(lt)
UpSet(cm, column_title = "Number of genes")
```

Next we look at the overlap of TSS with their exact positions.

```
lt = lapply(tss_lt, function(x) {
    unique(paste0(strand(x), seqnames(x), ":", start(x)))
})
cm = make_comb_mat(lt)
UpSet(cm, column_title = "Number of TSS (with their exact positions)")
```

```
tss_lt2 = lapply(tss_lt, function(x) {
    tb = table(x$gene_id)
    dp = names(tb[which(tb == 1)])
    x = x[x$gene_id %in% dp]
    names(x) = x$gene_id
    x
})
cn = tss_lt2[[1]]$gene_id
for(i in 2:length(tss_lt2)) {
    cn = intersect(cn, tss_lt2[[i]]$gene_id)
}
length(cn)
```

```
## [1] 17081
```

```
tss_lt2 = lapply(tss_lt2, function(x) x[cn])
```

```
library(GetoptLong)
compare_tss_pos = function(tss1, tss2, name1, name2, ...) {
    d1 = start(tss1)
    d2 = start(tss2)

    diff = abs(d1 - d2)
    
    v = numeric()
    v["0"] = sum(diff == 0)
    v["1-5"] = sum(diff >= 1 & diff <= 5)
    v["6-10"] = sum(diff >= 6 & diff <= 10)
    v["11-50"] = sum(diff >= 11 & diff <= 50)
    v["51-500"] = sum(diff >= 51 & diff <= 500)
    v["501-5kb"] = sum(diff >= 501 & diff <= 5000)
    v["5kb-50kb"] = sum(diff >= 5001 & diff <= 50000)
    v[">50kb"] = sum(diff >= 50001)

    barplot(v, ylab = "Number of TSS", 
        main = qq("TSS dist_diff, @{name1} and @{name2}\nmean (trim 0.05) = @{round(mean(diff, trim = 0.05))}bp, median = @{median(diff)}bp"), 
        las = 3, ...)
}

par(mfrow = c(4, 4))
for(i in 1:4) {
    for(j in 1:4) {
        if(i == j) {
            plot(c(0, 1), c(0, 1), type = "n", axes = FALSE, ann = FALSE)
            text(0.5, 0.5, names(tss_lt2)[i], cex = 1.5)
        } else {
            compare_tss_pos(tss_lt2[[j]], tss_lt2[[i]], names(tss_lt2)[j], names(tss_lt2)[i], ylim = c(0, 16000))
        }
    }
}
```

Top 10 TSS which the highest variability of their positions:

```
library(matrixStats)
pos_mat = do.call(cbind, lapply(tss_lt2, start))
v = rowSds(pos_mat)
ind = order(v, decreasing = TRUE)[1:10]
pos_mat2 = data.frame("chr" = as.vector(seqnames(tss_lt2[[1]])), pos_mat, Entrez_ID = tss_lt2[[1]]$gene_id)
pos_mat2 = pos_mat2[ind, ]

library(org.Hs.eg.db)
map = unlist(as.list(org.Hs.egSYMBOL))

pos_mat2$Entrez_ID = qq("[@{pos_mat2$Entrez_ID}](https://www.genecards.org/cgi-bin/carddisp.pl?gene=@{map[pos_mat2$Entrez_ID]}#genomic_location)", collapse = FALSE)
kable(pos_mat2, row.names = FALSE)
```

| chr | txdb\_known\_gene | gencode | refseq | great | Entrez\_ID |
| --- | --- | --- | --- | --- | --- |
| chr1 | 58546734 | 58546734 | 57424060 | 57424058 | 1600 |
| chrX | 37349330 | 38561542 | 38561542 | 38561370 | 7102 |
| chr11 | 41459773 | 41459773 | 41459652 | 40294115 | 57689 |
| chr3 | 75906695 | 75906695 | 77040099 | 75906695 | 6092 |
| chr8 | 31639222 | 31639222 | 32548311 | 32548635 | 3084 |
| chr17 | 34157294 | 34174964 | 33293295 | 33292989 | 40 |
| chr10 | 55627942 | 55627942 | 54801231 | 54801292 | 65217 |
| chr16 | 73891871 | 73891871 | 73048128 | 73131095 | 463 |
| chr16 | 5239802 | 5239802 | 6019024 | 6019703 | 54715 |
| chr3 | 24687919 | 24687887 | 25428263 | 25428311 | 5915 |

#### Influence on GREAT enrichment

TSS, although have different positions in different sources, are quite close. We next examnine whether the inconsistency of TSS positions affects the GREAT enrichment analysis.

In the next example, we use a dataset from UCSC table browser. The parameters are as follows:

```
clade = Mammal
genome = Human
assembly = GRCh38/hg38
group = Regulation
track = TF ChIP
table = A549 MYC (encTfChipPkENCFF542GMN)
```

Similarly, we perform local GREAT with five different TSS sources.

```
df = read.table("data/A549_MYC_encTfChipPkENCFF542GMN_hg38.bed")
df = df[df[, 1] %in% paste0("chr", c(1:22, "X", "Y")), ]
gr = GRanges(seqnames = df[, 1], ranges = IRanges(df[, 2]+1, df[, 3]))
res_txdb = great(gr, "GO:BP", "TxDb.Hsapiens.UCSC.hg38.knownGene", min_gene_set_size = 0)
res_gencode = great(gr, "GO:BP", "gencode_v40", min_gene_set_size = 0)
res_refseq = great(gr, "GO:BP", "refseq:hg38", min_gene_set_size = 0)
res_great = great(gr, "GO:BP", "great:hg38", min_gene_set_size = 0)

res_list = list(
    txdb_known_gene = res_txdb,
    gencode = res_gencode,
    refseq = res_refseq,
    great = res_great
)
```

We check the overlap of significant GO terms:

```
tb_list = lapply(res_list, function(x) getEnrichmentTable(x))

lt = lapply(tb_list, function(x) {
    x$id[x$p_adjust < 0.01]
})
cm = make_comb_mat(lt)
UpSet(cm, column_title = "Number of significant GO terms (FDR < 0.01)")
```

```
tb_list = lapply(res_list, function(x) getEnrichmentTable(x))
cn = intersect(tb_list[[1]]$id, intersect(tb_list[[2]]$id, intersect(tb_list[[3]]$id, tb_list[[4]]$id)))

vl = lapply(tb_list, function(x) {
    rownames(x) = x$id
    log2(x[cn, "fold_enrichment"])
})
par(mfrow = c(4, 4))
for(i in 1:4) {
    for(j in 1:4) {
        if(i == j) {
            plot(c(0, 1), c(0, 1), type = "n", axes = FALSE, ann = FALSE)
            text(0.5, 0.5, names(vl)[i], cex = 1.5)
        } else {
            plot(vl[[j]], vl[[i]], xlab = names(vl)[j], ylab = names(vl)[i], pch = 16, 
                col = "#00000020", main = "log2(Fold enrichment)",
                xlim = c(-6, 6), ylim = c(-6, 6))
        }
    }
}
```
